# Changes in Spo0A~P pulsing frequency control biofilm matrix deactivation

**DOI:** 10.1101/2025.02.13.638117

**Authors:** C.S.D. Palma, D.J. Haller, J.J. Tabor, O.A. Igoshin

**Author notes:** Corresponding author: Oleg Igoshin.

## Abstract

Under starvation conditions, *B. subtilis* survives by differentiating into one of two cell types: biofilm matrix-producing cells or sporulating cells. These two cell-differentiation pathways are activated by the same phosphorylated transcription factor - Spo0A~P. Despite sharing the activation mechanism, these cell fates are mutually exclusive at the single-cell level. This decision has been shown to be controlled by the effects of growth rate on gene dosage and protein dilution in the biofilm matrix production network. In this work, we explore an alternative mechanism of growth rate-mediated control of this cell fate decision. Namely, using deterministic and stochastic modeling, we investigate how the growth-rate-dependent pulsing dynamics of Spo0A~P affect biofilm matrix deactivation and activation. Specifically, we show that the Spo0A~P pulsing frequency tunes the biofilm matrix deactivation probability without affecting the probability of biofilm matrix activation. Interestingly, we found that DNA replication is the cell cycle stage that most substantially contributes to the deactivation of biofilm matrix production. Finally, we report that the deactivation of biofilm matrix production is not primarily regulated by the effects of growth rate on gene dosage and protein dilution. Instead, it is driven by changes in the pulsing period of Spo0A~P. In summary, our findings elucidate the mechanism governing biofilm deactivation during the late stages of starvation, thereby advancing our understanding of how bacterial networks interpret dynamic transcriptional regulatory signals to control stress-response pathways.

**Author Summary:** Bacteria have evolved various adaptation mechanisms to survive under challenging environmental conditions. For instance, under mild starvation, *B. subtilis* bacteria form biofilms — communities of cells encapsulated in a protective extracellular matrix. On the other hand, these bacteria differentiate into highly resistant spores under severe starvation. Interestingly, sporulating cells are often found within biofilm communities, but they do not contribute to biofilm matrix production. This is thought to be an energy-conservation strategy, as biofilm formation is an energy-intensive process and is therefore halted before sporulation begins. Though previous work has focused on the mechanisms driving biofilm disassembly, few studies have explored the regulatory processes that *B. subtilis* employs to halt matrix production prior to starting sporulation. In this study, we use mathematical models to demonstrate that the temporal dynamics of the biofilm master regulator Spo0A~P control the deactivation of matrix production. Understanding the regulation of biofilm, a common lifestyle in bacteria, can lead to the development of synthetic strategies to either enhance or disrupt biofilm formation, with potential applications in medicine and industry.

## 1. Introduction

Cell fate refers to the specific developmental path that a living cell takes, leading to its ultimate differentiation into a particular cell type with specific functions. Transcription factors can drive cell fate decisions by activating or repressing genes related to alternative fates (1,2). Understanding how transcription factors regulate cell-fate decisions is a fundamental question in biology because it determines how organisms maintain their functions and respond to the environment.

Bacteria including the soil microbe *B. subtilis* encounter challenging environmental conditions such as starvation. To ensure survival when facing starvation *B. subtilis* can differentiate into multiple distinct cell types. In response to mild starvation, individual bacterial cells differentiate from a motile to a sessile cell state (3). Cells in the sessile state produce and secrete an extracellular matrix, which leads to the formation of a biofilm (4). Biofilms are characterized by matrix-encased communities of surface-associated cells that provide resistance to stresses such as antibiotics (4,5). In contrast, prolonged starvation induces sporulation, wherein cells differentiate into metabolically-inactive spores that resist heat, chemical toxins, and other stressors (6–9).

The decision to form a biofilm or sporulate is governed by gene regulatory networks that are activated by the same major regulatory transcription factor, named Spo0A (0A) (10,11). The concentration of 0A is controlled transcriptionally (12) and phosphorylation governs its activity post-translationally (2,13,14). The phosphorylated form of 0A (0A~P) induces expression of sporulation and biofilm genes (2). The phosphorylation of 0A is controlled by a phosphotransfer cascade (Figure 1A) termed phosphorelay (14). The phosphorelay cascade, initiates with five histidine kinases (KinA, KinB, KinC, KinD, and KinE) that autophosphorylate (15,16), with KinA being the major sporulation kinase (17). Next, the phosphate from each of the kinases is transferred to two phosphotransferases Spo0F (0F) and Spo0B (0B), and finally to 0A (14,18). Further, 0A~P concentration is also controlled by the phosphatases Spo0E and Rap, which dephosphorylates 0A~P and 0F~P, respectively (19).

**Figure 1:**
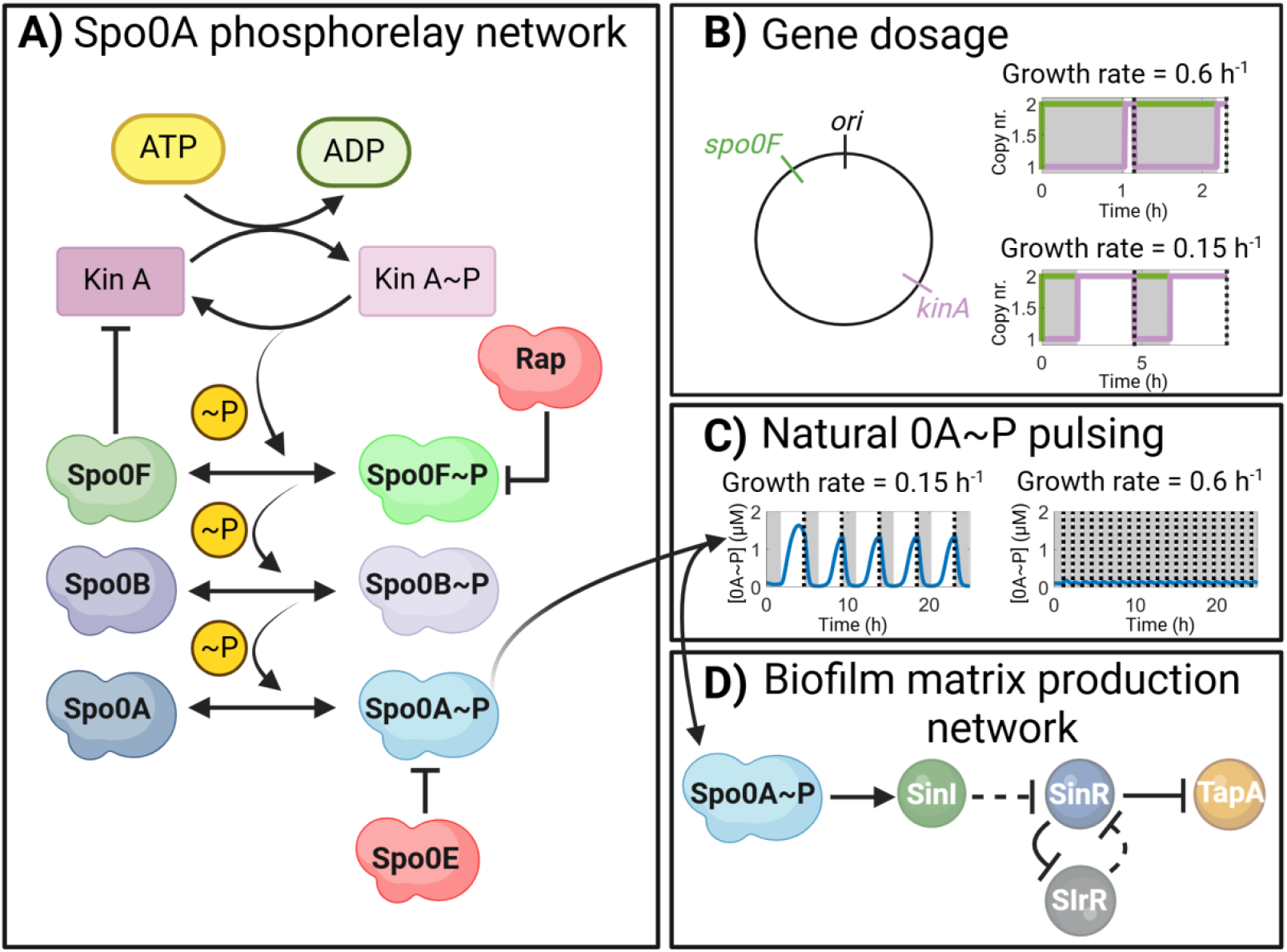
Illustration depicting the networks analyzed. **A)** Post-translational phosphorelay network of Spo0A (0A) (14). Kinase A (KinA) transfers phosphoryl groups to Spo0A via the two phosphotransferases Spo0B and Spo0F (18). Phosphorylated Spo0F and Spo0A are subject to negative regulation by phosphatases Rap and Spo0E, respectively (19). Spo0F inhibits KinA activity (20). **B)** Illustration of gene dosage imbalance for *kinA* and *spo0F*. Shown are the relative positions of *kinA (126° - terminus* proximal) and *spo0F (326° - oriC* proximal) in the genome (33). Also shown are the approximate gene copy numbers over two cell cycles for a growth rate of 0.6 h^−1^ (upper plot) and a growth rate of 0.15 h^−1^ (bottom plot). Grey areas correspond to the DNA replication time, estimated using *(Eq. 11)*. Dashed lines correspond to end of the cell cycle, estimated from *(Eq. 12)*. **C)** Predicted Spo0A~P natural pulsing dynamics by the phosphorelay network model assuming a growth rate equal to 0.15 h^−1^. For comparison, also shown is the 0A~P level assuming a growth rate equal to 0.6 h^−1^. **D)** Biofilm matrix production network. Spo0A~P induces the production of SinI (23,24). SinI represses the activity of SinR. SlrR and SinR form a double negative mutual repressive network (28,34). Dashed arrows depict regulatory effects due to post-translational interactions i.e., the activity of SinR is regulated by complex formation with SinI and SlrR (28,34). When SinR is in complex with SinI or SlrR, the expression of *tapA* and *slrR* are de-repressed. Figure created with BioRender.

Previously, we demonstrated that 0A~P has pulsatile dynamics driven by a transient gene dosage imbalance and a negative feedback loop in the phosphorelay network (17). Specifically, 0F inhibits KinA activity and the chromosomal arrangement of *kinA* (terminus-proximal) and *spo0F* (*oriC*-proximal) (Figure 1B) leads to gene dosage imbalance during the DNA replication period. Since *spo0F* is replicated at the beginning of replication while *kinA* is replicated near the end of the replication period, the DNA replication time is about the same length as the cell cycle during high growth rates (17), leading to a gene dosage ratio *kinA*:*0F* below 1:2 (upper plot in Figure 1B). These conditions lead to overall low levels of 0A~P in the cell (Figure 1C, right plot). Conversely, for low growth rates, the cell cycle is longer than the DNA replication time (17), leading to a balanced gene dosage ratio of 2:2, after DNA replication (bottom plot in Figure 1 B). This leads to low 0A~P levels during the DNA replication period, followed by an overshoot after the replication period and prior to the cell division. In addition, the growth slowdown leads to an increase in the concentration of KinA which further contributes to the overshoot of 0A~P amplitude (20). This once-per-cell-cycle overshoot response leads to the pulsatile dynamics of 0A~P (21) (Figure 1C, left plot).

Once phosphorylated, 0A can activate the matrix production genes via the biofilm matrix production network (Figure 1D)(11,22). Namely, 0A~P induces the expression of transcription factor SinI which is known to inactivate the transcription factor SinR by sequestration in a complex (23–26). Active SinR inhibits the expression of SlrR, which can also inactivate SinR by sequestration (27,28). Inactivation of SinR results in the derepression of a set of genes and operons controlling biofilm formation, including the *tapA* gene that encodes a protein required for biofilm matrix formation (28–30).

On the other hand, 0A~P is known to directly activate sporulation genes (31). The activation of genes involved in sporulation requires a higher threshold of 0A~P than biofilm matrix production genes (2). However, biofilm and sporulation as cellular fates are mutually exclusive (i.e., cells do not activate them simultaneously). Therefore, there should exist regulatory mechanisms that ensure biofilm matrix production deactivation when 0A~P levels exceed the threshold to trigger sporulation.

Recently, it was suggested that the biofilm matrix deactivation under starvation can be explained by the slowdown in growth rate (32). In particular, the biofilm matrix production network was found to be bistable for 0A~P levels higher than a certain threshold. As growth slows down, there will be an insufficient amount of SinI and SlrR to fully inhibit the activity of SinR. As a result, the system enters a monostable region where matrix production is not active, i.e., *tapA* expression is low. However, that study focused on the population-level average concentration of 0A~P. Consequently, the impact of 0A~P pulsatile dynamics on the deactivation of biofilm matrix production remains unexplored at the single-cell level.

Here we build on previous mathematical models (20,32) to predict how the biofilm matrix production network decodes 0A~P pulsing signals to regulate biofilm matrix production. Specifically, we examine how the frequency and amplitude of the pulsatile signal influence the biofilm matrix production network and, consequently, the cell-fate decision in *B. subtilis*.

## 2. Results

### 2.1 Biofilm is deactivated for a certain frequency and amplitude of the 0A~P natural pulsing signal

To investigate the effects 0A~P pulsing has on biofilm matrix deactivation, we coupled a deterministic phosphorelay network model (Methods Section 4.1) (20) with a deterministic biofilm matrix production network model (32) (Methods Section 4.2) and performed a deterministic simulation assuming decreasing growth rate (Figure 2A). We began by comparing the effects of 0A~P pulsing dynamics with those of a constant 0A~P level, corresponding to the average of the pulsing signal, across each cell cycle (Figure 2B). Starting from a matrix active state, our results show that for constant 0A~P levels, the production of the matrix protein TapA is not deactivated. However, assuming 0A~P pulsing, TapA production is deactivated earlier in time, i.e., at a faster growth rate. (Figure 2C). Overall, this suggests that the frequency and amplitude of 0A~P natural pulsing may act as a regulator of biofilm matrix deactivation. As such, not accounting for the pulsatile behavior of 0A~P can lead to incorrect conclusions about the mechanisms controlling cell fate decisions in *B. subtilis*.

**Figure 2:**
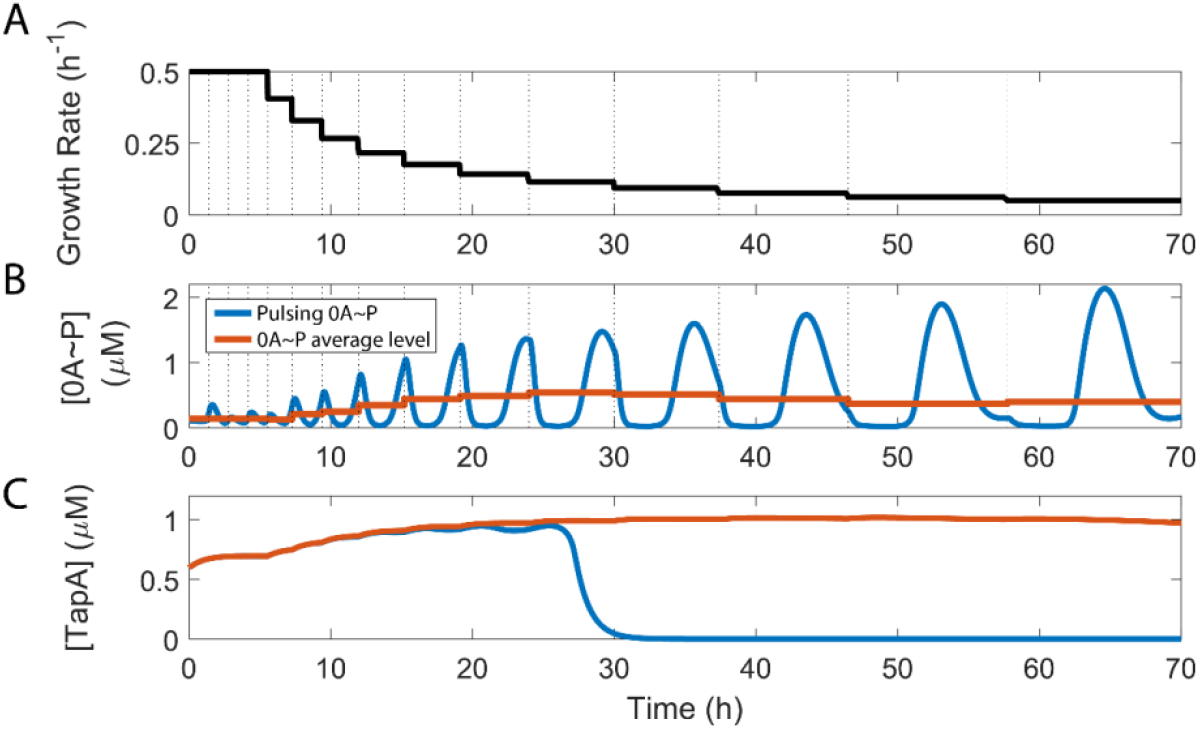
Effects of 0A~P pulsing dynamics control biofilm matrix deactivation. **(A)** Growth rate dynamics used as the input of the phosphorelay and biofilm matrix production mathematical models **(B)** 0A~P predicted levels from the phosphorelay model assuming the growth rate input dynamics shown in A (blue line). Also shown is the 0A~P pulse average level per cell cycle (orange line). Vertical dashed lines represent the beginning of a new cell cycle, according to Eq.12. **(C)** Predicted levels of biofilm matrix protein TapA, estimated from the biofilm matrix production network model. Blue line corresponds to the TapA predicted levels assuming the 0A~P pulsatile input shown in blue color in B. Orange line corresponds to the TapA predicted levels assuming the constant level of 0A~P, per cell cycle, shown by the orange line in B.

### 2.2 Pulsing dynamics of 0A~P affect the probability of matrix deactivation but not activation

In single cells, matrix activation and deactivation are stochastic processes (32). To examine how these processes depend on the 0A~P pulsing dynamics, we developed a stochastic model of the biofilm matrix production network, based on (32) (Methods Sections 4.2 and 4.3).

We performed stochastic simulations (Methods Section 4.4) using the natural pulsing 0A~P dynamics assuming a growth rate equal to 0.4 h^−1^ and the corresponding average signal (Figure 3A) as inputs. For each of the inputs, we simulated over 2000 cells for a total of 40 hours, starting with cells in both a matrix-inactive state (OFF, Figure 3B) and a matrix-active state (ON, Figure 3C), at time = 0 h. We note that to model the cells’ behavior accurately, each cell state at t = 0 h was set to be the final state of a 60 h preliminary stochastic simulation. The results show that, independently of the initial cell state, both input types can activate or deactivate the production of matrix protein TapA (Figure 3B-C).

**Figure 3:**
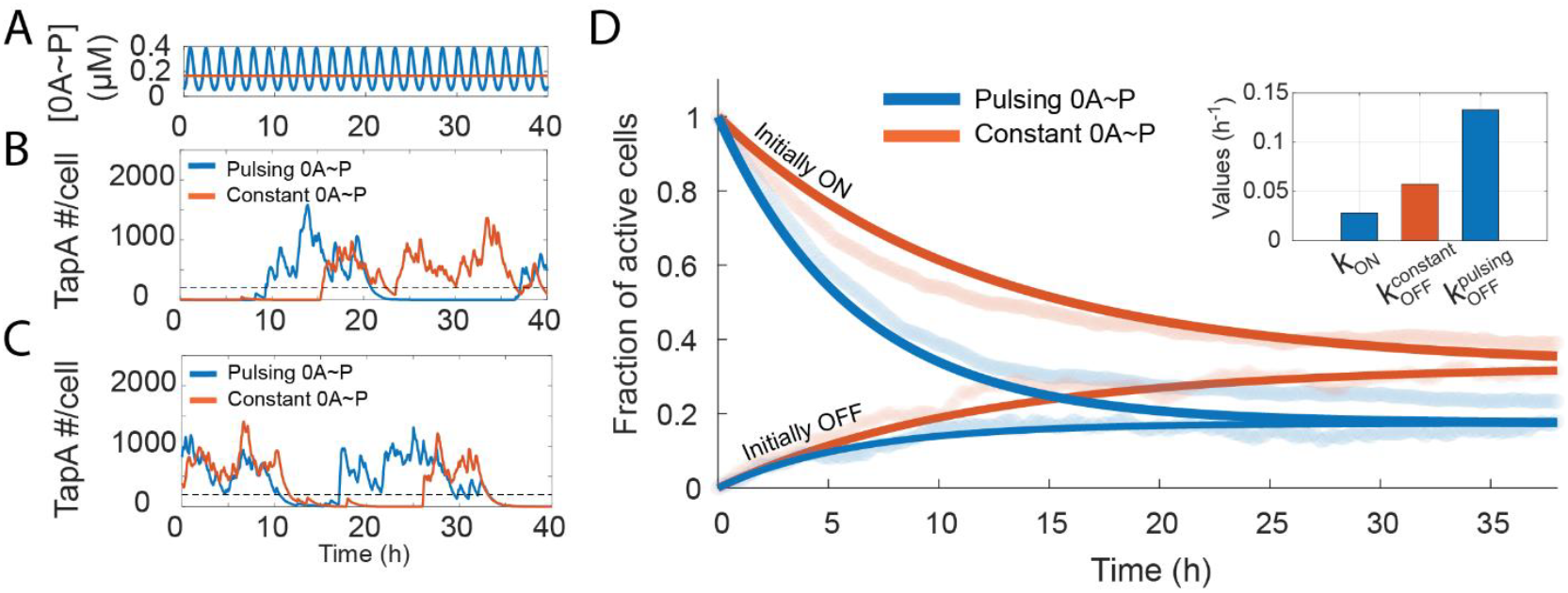
0A~P pulsing tunes the biofilm matrix deactivation rate. **(A)** Pulsing (blue) and constant (orange) 0A~P signal used as input to the biofilm matrix production stochastic model. Pulsing signal corresponds to the signal for growth rate equal to 0.4 h^−1^ predicted by the phosphorelay network model. The constant 0A~P signal corresponds to the mean of the pulsing signal. **(B)** Stochastic simulations starting from biofilm matrix inactive state for both constant and pulsing 0A~P signal. Matrix production is considered inactive if the number of TapA molecules < 200 (dashed line). **(C)** Stochastic simulations starting from matrix active state for both constant and pulsing 0A~P signal. Matrix production is considered active if the number of TapA molecules ≥ 200 (dashed line). **(D)** Fraction of active cells as a function of time for constant (dark orange) and pulsing 0A~P signal (dark blue) estimated from fitting *Eq. 3* and *Eq. 4* to the stochastic simulation data of ‘*Initially OFF cells*’ (light blue) and ‘*Initially ON’* (light orange) cells, respectively. All fits have R^2^ >= 0.77 and MSE < 0.005. A total of 2000 simulations were performed. **(inset)** Estimated values of the biofilm activation rate (*k*_*ON*_) and of the matrix deactivation rate assuming pulsing 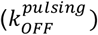 and constant 0A~P input 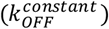.

Next, to investigate the effects of 0A~P pulsatile dynamics on matrix activation we calculated the probability of matrix activation and deactivation over time. Namely, from the stochastic simulation results, at each time point we calculated the fraction of cells with matrix active (*F*^*ON*^):

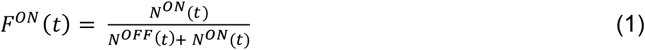

Here *N*^*ON*/*OFF*^ (*t*) is the number of cells with matrix production active/inactive, respectively, at time *t*. We considered a TapA threshold ≥ 200 molecules to be associated with biofilm matrix production. This threshold was determined based on the bimodal distribution of TapA molecules (Supplementary Figure S2). The fraction of active cells over the course of the stochastic simulations are shown in light orange and light blue color in Figure 3D.

To understand the observed dynamics of *F*^*ON*^ we employ a simple two-state model *ON* ↔ *OFF* that assumes a Markov process (e.g., memory-less activation and deactivation) with effective rates *k*_*ON*_ and *k*_*OFF*_. For this model, we can write the probability of being in the active (ON) state as:

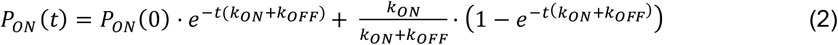

For *P*_*ON*_ (0) = 0 (*i*.*e*., all cells have matrix production initially OFF), *Eq. 2* becomes:

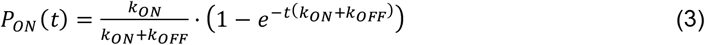

Meanwhile, for *P*_*ON*_ (0) = 1 (*i*.*e*., all cells have matrix production initially ON), *Eq. 2* becomes:

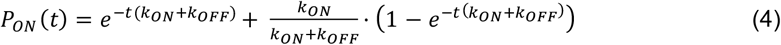

At steady state (*i*.*e*., as *t* → ∞) both *Eq. 3* and *Eq. 4* approach the same steady state fraction of active cells:

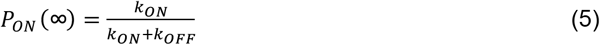

On the other hand, at short times, *Eq. 3* can be approximated by a polynomial expression using a Taylor series expansion:

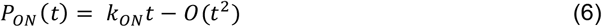

Therefore, initially the fraction of active cells linearly increases with a slope given by *k*_*ON*_ :

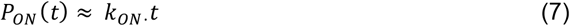

Following the same rationale, approximation of *Eq. 4* by Taylor series expansion yields that the linear decrease of activation probability for the case when all cells are initially in an ON state:

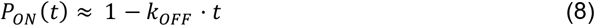

Therefore, by comparing activation dynamics in Figure 3D ‘Initially OFF’ (light blue and orange lines), one can see that both have approximately the same initial slope (for t ⪅ 2.5 h). According to *Eq. 7*, this suggests that both constant and pulsatile dynamics of 0A~P have the same matrix activation rate, *k*_*ON*_· On the other hand, the difference in the slopes of deactivation kinetics (blue and orange lines of ‘Initially ON’ cells) suggests, according to *Eq. 8*, distinct matrix deactivation rates, *k*_*OFF*_, for constant and pulsatile dynamics of 0A~P.

To estimate *k*_*ON*_ and *k*_*OFF*_ rates based on the whole range of simulated times, we simultaneously fit *Eq. 3* to ‘‘*Initially OFF*’ cells and *Eq. 4* to ‘*Initially ON*’ cells (Methods Section 4.6). Fitting results (dark orange and blue lines in Figure 3D) yield a good fitting (MSE < 0.005) when imposing a single *k*_*ON*_, for both pulsing and constant 0A~P input. Meanwhile, as expected, *k*_*OFF*_ estimation for pulsing input is higher than for constant 0A~P input (inset Figure 3D). Overall, our results show that pulsing dynamics of 0A~P do not affect matrix activation probability but increase the matrix deactivation probability.

### 2.3 Period of 0A~P oscillation controls the biofilm matrix deactivation threshold

Reference (32) investigated the bistability of the biofilm matrix production network assuming a constant level of 0A~P. Namely, for a given concentration of 0A~P, a deterministic model of the network predicts the existence of two stable steady states of TapA levels. For a 0A~P concentration lower than ~0.1 µM the system is monostable with only a matrix OFF steady state (Figure 4A). For a 0A~P concentration higher than ~0.1 µM the system is bistable with an OFF and an ON matrix stable steady-state of TapA (Figure 4A, grey area).

**Figure 4:**
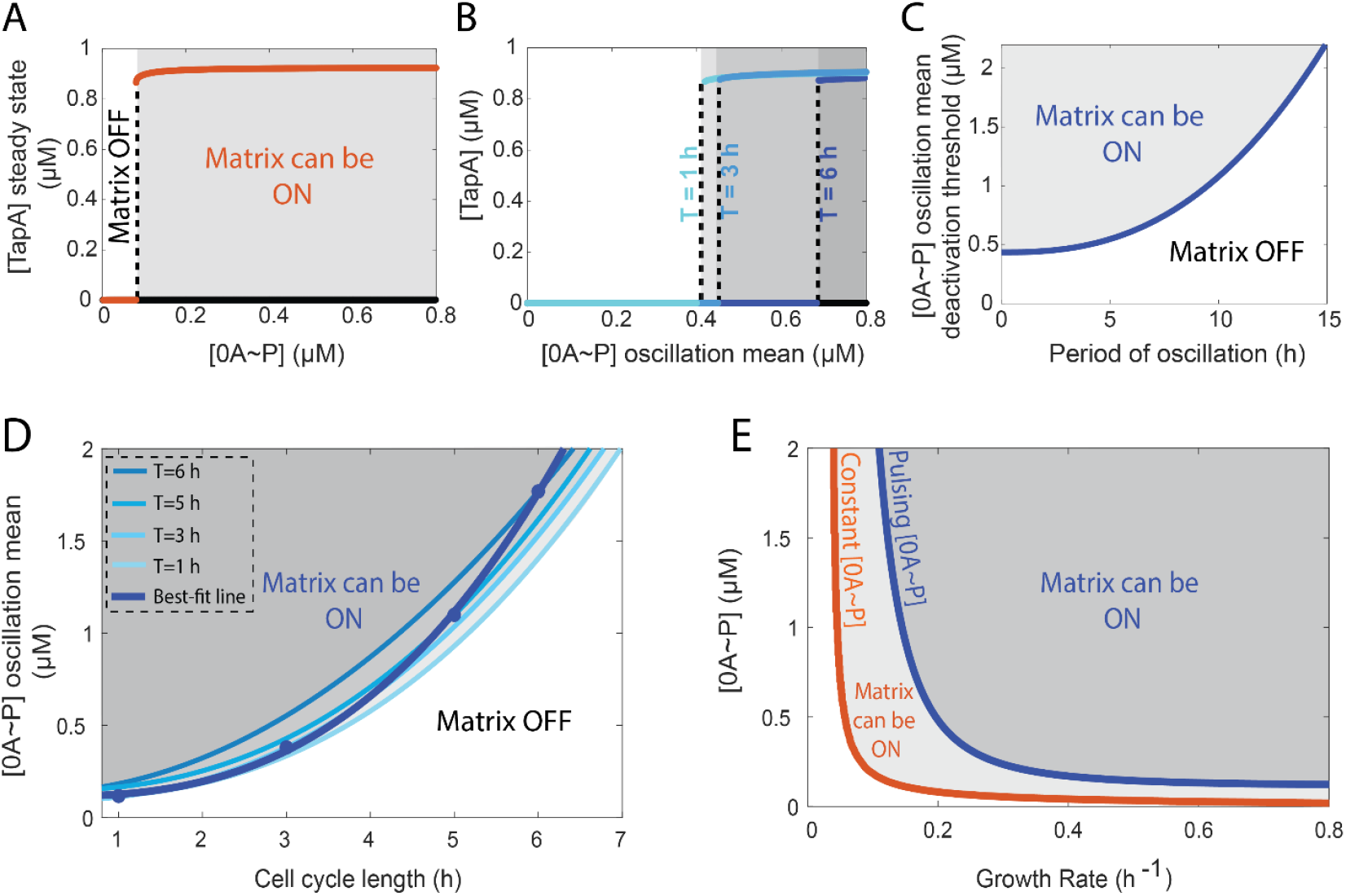
Biofilm matrix deactivation threshold increases with increasing 0A~P oscillation period. **(A)** Bistability diagram of the biofilm matrix production network assuming a constant level of 0A~P. Dashed line represents the concentration of 0A~P that corresponds to the biofilm matrix deactivation threshold. To the right of the dashed line the system presents bistability where two steady states co-exist: one corresponding to high levels of matrix protein TapA and the other corresponding to deactivated biofilm matrix production (i.e. low levels of TapA). For 0A~P concentrations below the threshold the system is monostable with low levels of TapA production (matrix production OFF). The results shown are for a growth rate of 0.2 h^−1^. **(B)** Biofilm deactivation threshold as a function of the 0A~P oscillation mean for three oscillatory signals of different periods. Oscillations were programmed as cosine functions of the form [0*A*~*P*] = *amplitude* · *cos* (*freq. t*) + *offset*. The oscillation periods tested were 1, 3 and 6 h which correspond to an angular frequency (*freq*) of ~6.28 h^−1^, 2.09 h^−1^, and 1.05 h^−1^, respectively. The *offset* was set such that the minimum value of the [0A~P] oscillation is 0. The results shown are for a growth rate of 0.2 h^−1^. **(C)** [0A~P] biofilm deactivation threshold (i.e. the 0A~P oscillation mean amplitude) as a function of the period of oscillation. **(D)** Bifurcation between regions for which matrix can be active (ON) and inactive (OFF) as a function of the cell cycle length assuming 0A~P oscillatory signals of different periods (T). Shown is the bifurcation deactivation threshold for T=1, 3, 5, and 6 h (light blue lines). Also shown in the best fit power-law (Methods Section 4.6) estimated from the datapoints for which the oscillation period matches the cell cycle length (dark blue line). **(E)** Bifurcation between regions for which matrix can be active (ON) and inactive (OFF) as a function of growth rate. Orange line shows the results assuming a constant level of 0A~P. Blue line shows the result assuming a period of oscillation of 0A~P equal to the cell cycle length (estimated from dark blue line in D).

To examine the effects of 0A~P pulsing dynamics on the biofilm deactivation threshold (i.e., the average 0A~P concentration for which the system shifts from a bistable to a monostable OFF state), represented by the dashed line in Figure 4A, we generated *in silico* 0A~P oscillatory signals with varying amplitudes and periods. This approach allowed us to investigate how these parameters influence the deactivation threshold by decoupling the effects of amplitude and period from the natural pulsing behavior of 0A~P. In Figure 4B we show the effects that three different 0A~P oscillatory input signals have on the deactivation threshold. Specifically, periods (T) of 1, 3, and 6 hours were tested. The results show that as the period increases (*i*.*e*., the frequency decreases), the matrix deactivation threshold also increases. This implies that a longer oscillation period requires a higher amplitude of the 0A~P signal to maintain the matrix production (ON) state. Notably, the threshold is almost unresponsive to changes in period for short oscillation periods but becomes highly responsive to longer periods (Figure 4C). This indicates that the system deactivation threshold exhibits varying degrees of responsiveness depending on the oscillation period. Altogether, these results show that as the period increases the amplitude of the 0A~P oscillatory signal must increase significantly to sustain biofilm matrix production.

Next, we examined how the deactivation threshold is affected when there is one 0A~P pulse per cell cycle, *i*.*e*., situation reflecting natural conditions (17). For this study, we began by analyzing how the deactivation threshold varies with the cell cycle length for oscillatory periods of 1, 3, 5 and 6 hours (light blue lines in Figure 4D). Next, for each line, we selected the value for which the oscillation period matches the cell cycle length and fit a power-law (Methods Section 4.6). The best fit line (dark blue line in Figure 4D) shows that the biofilm deactivation threshold increases as a function of the cell cycle length if one pulse occurs per cell cycle.

Finally, to examine the effects of pulsing versus a 0A~P constant signal on matrix deactivation across different growth rates, we compare the deactivation threshold as a function of the growth rate for an oscillatory signal occurring once per cell cycle (estimated from the best-fit line in Figure 4D) with that of a constant 0A~P signal. For both types of signals, the results (Figure 4E) show that as cells grow slower, the required level of 0A~P to keep the matrix active (ON) becomes higher. However, the effects of oscillatory dynamics significantly affect when the matrix production is deactivated, causing it to occur earlier (i.e., higher growth rates) compared to constant 0A~P dynamics.

### 2.4 DNA replication time increase is the determinant factor for biofilm deactivation

Given that the natural 0A~P pulsing period matches the cell cycle duration (Figure 1B) and matrix deactivation occurs at a certain cell cycle length, we investigated which phase of the cell cycle (DNA replication or post-replication) is responsible for controlling matrix deactivation.

We started by building a simple model where the 0A~P input is set to be a square pulsing signal. We tested two different square pulsing signals. For the first signal the 0A~P ON time is kept constant, and the OFF time increases in each cycle (Figure S1A). For the second signal the opposite occurs (Figure S1B). Next, we tested the effects of these signals on matrix deactivation. We found that an increase in OFF time drives matrix deactivation, whereas increase in ON time does not (Figure S1C).

In the natural 0A~P pulsing signal, the equivalent of the OFF-time period is the DNA replication period since the gene dosage ratio *kinA*:*0F* is approximately 1:2 during this period, resulting in low levels of 0A~P (17). To test if the increase in the DNA replication period is the major regulator of biofilm matrix deactivation, we performed a similar test as the one for the simple model. Specifically, we simulated the phosphorelay network model under two different scenarios of cell-cycle slowdown (Methods Section 4.4). In the first one, the cell cycle and the DNA replication time increase by the same amount and the post-replication period is kept constant (Figure 5A). In the second one, the cell cycle increases together with the post-replication period, but DNA replication period is kept constant (Figure 5B). For each scenario, we tested the effects of the increase in the cell cycle on the deactivation of biofilm matrix production. We found that the biofilm matrix deactivates at shorter cell cycles for the network in which the DNA replication period increases, and the post-replication period is kept constant (dark blue bar in Figure 5C). However, unlike the simple model, we also observe deactivation for the 0A~P signal when only post-replication time increases (light blue bar in Figure 5C). This result suggests that post-replication time also plays a regulatory role in matrix deactivation, but its influence is smaller than that of the DNA replication period.

**Figure 5:**
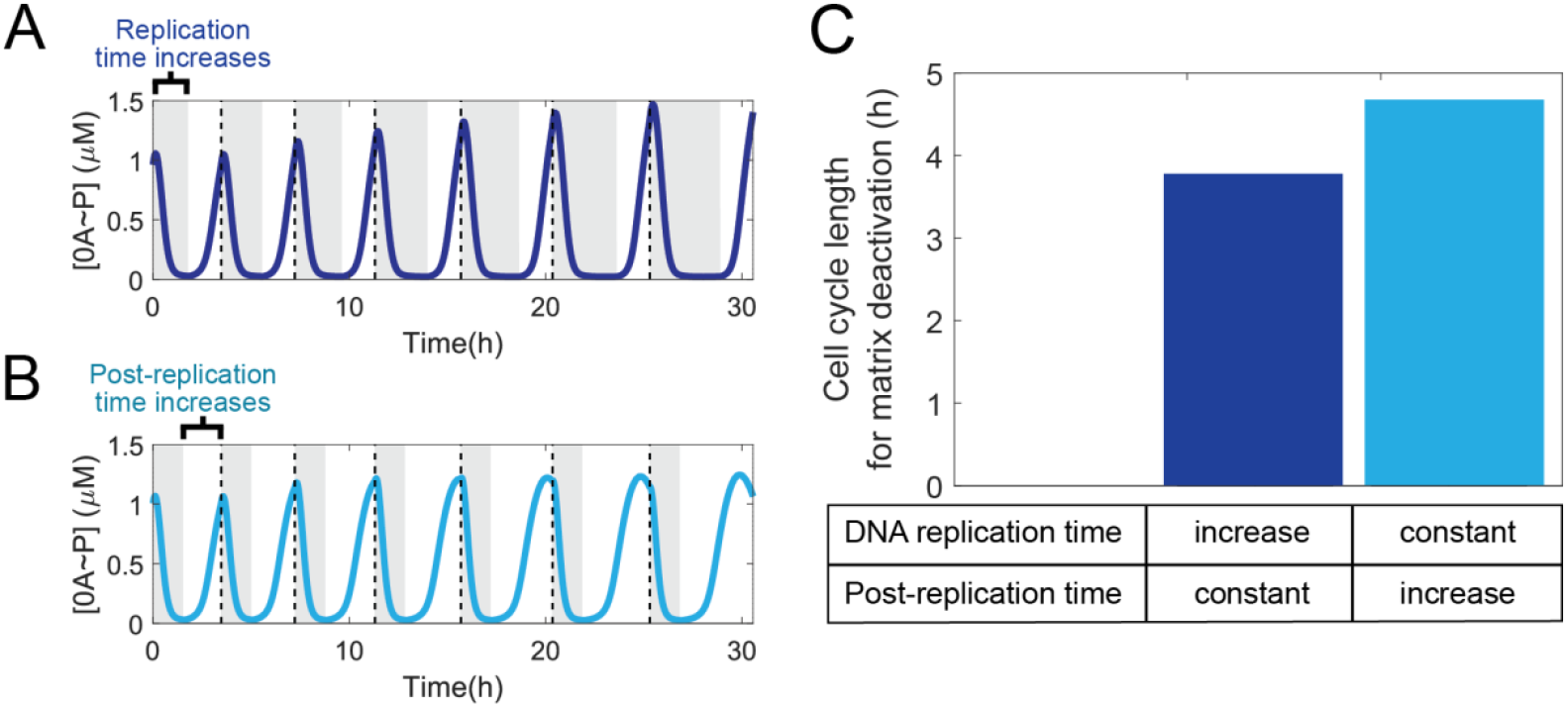
DNA replication time is the cell-cycle stage that mostly contributes to biofilm matrix deactivation. **(A)** 0A~P pulsing dynamics for increasing DNA replication period (grey regions) and constant post-replication period (white regions). Pulsing occurs once per cell-cycle. Dashed lines represent the end of a cell cycle. **(B)** The same analysis as (A) for constant DNA replication period and increasing post-replication period. **(C)** Cell cycle length for which biofilm matrix is deactivated when cell cycle time increases by solely increasing DNA replication period (dark blue bar) or the post-replication period (light blue bar).

### 2.5 The growth rate influences biofilm deactivation through multiple mechanisms, with the contribution of the 0A~P pulsing period being the most significant

Previous research identified the growth effects on the biofilm network, namely the effects on gene dosage and protein dilution, as the primary regulators of biofilm matrix deactivation (32). Here, we compared the impact of such effects to that of changes in 0A~P pulsatile signal dynamics in controlling biofilm matrix deactivation.

We began by simulating the biofilm matrix production network for five different conditions (Methods Section 4.6). Each condition differs in whether it includes the growth rate effects on gene dosage, protein dilution, and/or the 0A~P pulsatile signal. Specifically, starting with an initial condition where biofilm matrix production is active (set at a growth rate of 0.2 h^−1^), we simulated the network, using 0A~P natural pulsing as input, to determine the minimum cell cycle length leading to biofilm matrix production deactivation (hereafter ‘cell cycle deactivation threshold’). Specifically, we ran deterministic simulations for increasing cell cycle lengths until reaching steady-state concentration of the model species (>100 h). Our results (black bar in Figure 6) show that the cell cycle deactivation threshold is 3.9 h. Next, for condition two, we examine how growth rate-mediated changes in protein dilution impact the cell cycle deactivation threshold. To this end, we simulated the network for effective protein degradation rates (*Eq. 18*) without growth rate dependence (*Eq. 19*). Under these conditions, matrix production deactivation occurs at a slightly longer cell cycle deactivation threshold, equal to 4 h. Similarly, in condition three, to examine the impact of gene dosage growth rate effects on the deactivation cell cycle threshold, we modified the average gene copy number (*Eq. 15*) to be independent of growth rate (*Eq. 20)*. For this condition, matrix deactivation occurred at a cell cycle threshold equal to 4.4 h. For condition four, we simulated the network with the simultaneous removal of both these effects, which extended the deactivation cell cycle threshold to 4.8 hours (Figure 6). Thus, the effects of growth rate on gene dosage and protein dilution in the biofilm matrix production network change the deactivation threshold by less than 23%. In contrast, a much larger effect is observed for the final tested condition that assesses the impact of 0A~P pulsing dynamics, while maintaining all other growth rate effects. When we replaced the pulsatile 0A~P signal at a growth rate of 0.2 h^−1^ with a constant 0A~P signal equivalent to the average value of the pulsatile signal, we observed an approximate 3-fold increase in the deactivation cell cycle threshold (Figure 6). We conclude that the 0A~P pulsatile signal is the primary regulator of biofilm matrix deactivation, outweighing the previously identified effects of growth rate on gene dosage and protein dilution.

**Figure 6:**
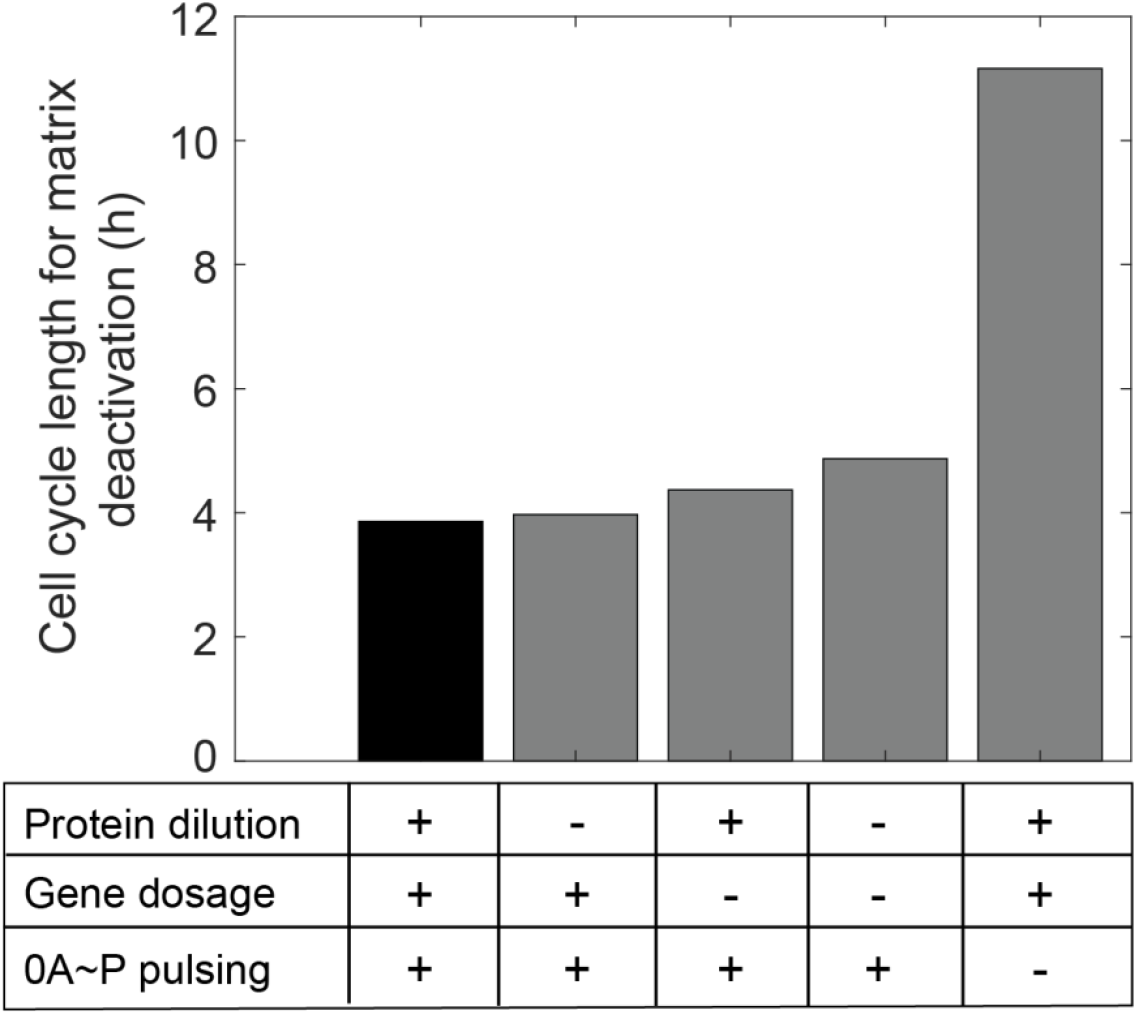
0A~P pulsing is the primary regulator of the minimum cell cycle length leading to biofilm matrix production deactivation. Shown is the cell cycle length for which biofilm matrix is deactivated for five different networks differing in the presence (+) or absence (−) of growth rate effects on protein dilution, gene dosage and/or 0A~P pulsing. Black bar indicates the result for the complete network with all growth rate effects present, assuming a growth rate equal to 0.2 h^−1^.

## 3. Discussion

Multiple single-cell phenotypes coexist in *B. subtilis* biofilms (6,33,34). For example, spore-producing cells are often found near the aerial boundary of the biofilm (22). It has been demonstrated that sporulating cells halt biofilm matrix production (35), likely because matrix production is an energy-intensive process requiring macromolecule synthesis, leading cells to reallocate energy towards sporulation (36–38). To successfully achieve this outcome, the decision to initiate matrix production and later halt it and transition to sporulation must be tightly regulated. The gene regulatory networks controlling the initiation of matrix production have been extensively studied (3,6,38), highlighting the importance of precise transcriptional control in the synthesis of matrix components. In contrast, the mechanisms regulating the transition from a matrix production to a sporulating phenotype are less understood and subject of debate. To date, few transcriptional control mechanisms have been proposed. First, high levels of Spo0A~P negatively regulate *sinI* expression leading to repression of matrix production (35). Second, changes in the gene dosage between *sinR* and *slrR* during sporulation are believed to repress matrix production (35). However, recent studies showed that artificially induced high 0A~P levels can fail to repress matrix production genes (15), and instead, a slowdown in growth rate plays a critical role in altering the dosage of *sinR* and *slrR*, leading to the repression of matrix production (32). Here, we describe another mechanism, based on the natural pulsatile dynamics of Spo0A~P, which has a stronger regulatory effect on the transition between matrix production and sporulation than all previously identified mechanisms. This mechanism not only enhances our understanding of matrix deactivation but also sheds light on the regulation of mutual exclusivity between biofilm and sporulation phenotypes.

Our results suggest that the period of 0A~P pulses, determined by the length of the cell cycle, serves as the primary regulator of biofilm deactivation. Specifically, as the growth rate decreases (i.e., as the cell cycle lengthens), both the period and amplitude of 0A~P pulses increase (Figure 2A and B). However, for long cell cycles (i.e. low growth rates), maintaining biofilm activity requires an exceptionally high increase in 0A~P pulse amplitude (Figure 4C). Consequently, while the 0A~P pulse amplitude does increase, it fails to reach the threshold necessary to sustain biofilm matrix production. As a result, biofilm is deactivated. Nevertheless, the amplitude of the 0A~P signal has been shown to reach the threshold necessary to trigger sporulation (20). Thus, we can conclude that the pulsing period controls biofilm deactivation whereas the pulsing amplitude triggers sporulation. This mechanism provides further insight into how the mutual exclusivity of these two phenotypes is maintained.

Every cell cycle, the completion of DNA replication triggers a 0A~P pulse, which in turn serves as the activation signal for sporulation (17), thereby ensuring that sporulation is not triggered prior to DNA replication completion. Here we report an additional feature controlled by the DNA replication period. Namely, beyond ensuring that sporulation occurs at the appropriate time, DNA replication is also the cell-cycle stage mostly responsible for the biofilm matrix production deactivation (Figure 5C). This mechanism ensures that under rapid decrease of growth rate cells deactivate matrix production during the DNA replication period before peak 0A~P induces sporulation. As such, we propose that, during each cell cycle cells reassess, whether to continue biofilm production or cease it and initiate sporulation.

Interestingly, although the pulsing signal reaches higher levels than the corresponding average signal, we did not find evidence that 0A~P pulsing dynamics significantly affect the activation of biofilm matrix production (Figure 3). Therefore, we conclude that activation is mostly controlled by the amplitude modulation of the 0A~P signal whereas deactivation is controlled by the frequency modulation of the signal. Notably, frequency modulation has previously been reported for the alternative sigma factor, σ^B^. Specifically, σ^B^ regulates the *B. subtilis* stress response by controlling its target genes through sustained pulsing, where higher stress levels are encoded by increased pulse frequency. Altogether, this indicates that pulse modulation may be a widespread regulatory mechanism in bacteria (39).

While the models used in this study have been experimental validated in prior research (20,32), we lack experimental validation of the results reported here. The major limitation to such validation is the absence of methods capable of precise dynamic control of transcription factors. Existing approaches, such as static gene perturbations (e.g., knockouts) (40) and basic dynamic control using chemical inducers (2), are not efficient for studying the fast natural temporal dynamics of transcription factors. Therefore, new approaches, for example, based on optogenetics or microfluidics (41), must be developed to enable precise, synthetic fine-tuning of transcription factor dynamics, to facilitate the *in vivo* study of how gene regulatory networks respond to such signals.

Overall, our results elucidate how *B. subtilis* utilize dynamical 0A~P signaling to drive distinct stress-response phenotypes. Interestingly, mechanisms wherein TF activity patterns dictate distinct phenotypes have been reported for eukaryotes (42). To our knowledge, our study is the first to report a similar mechanism in bacteria. Given that the GRN controlling biofilm and sporulation are widely conserved in pathogenic bacterial species such as *B. anthracis* (43) and *B. cereus* (44), we expect our findings to have implications in understanding the survival mechanisms of these bacteria, which is critical to understand their ability to cause infection. Ultimately, this study aims to contribute to the study of dynamic transcription factor activity into bacteria research on cell fate decision, moving beyond the traditional steady-state approach.

## 4. Methods

### 4.1 Phosphorelay model

We extended a previous mathematical model of the 0A phosphorelay (20) to uncover the effects of 0A~P pulsing in the cell fate decision of *B. subtilis*. The reactions and parameters used in this model are listed in Table S1. The production of the phosphorelay proteins and phosphatases (KinA, Spo0F, Spo0B, Spo0A, Rap, and Spo0E) is modeled by reactions R1 to R6. Post-translation interactions of phosphorylation/dephosphorylation are modeled by reactions R7 to R13. Specifically, after phosphorylation of the major sporulation kinase KinA (R7) the phosphoryl group is transferred to Spo0F (R8). Next, the phosphoryl group is transferred to Spo0B (R10) and finally to Spo0A (R11) (10,40). Phosphorylated Spo0F and Spo0A are subject to negative regulation by phosphatases Rap (R12) and Spo0E (R13), respectively. Reaction (R9) accounts for the effect of inhibition of KinA by Spo0F.

The expression of phosphorelay proteins is modeled with Hill-functions. To account for the delay induced by indirect feedbacks on the production of the phosphorelay proteins we assumed that 0A~P induces the expression of an intermediate regulator S, as in (20). The intermediate regulator induces the expression of *kinA, spo0F* and *spo0A* in accordance with the following generic Hill-function:

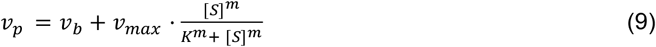

Where, *ν*_*b*_ is the basal rate, *ν*_*max*_ is the maximum expression rate when promoter is fully induced. The variables *K* and *m* are the half-maximal binding constant and the Hill-exponent, respectively. The variable [*S*] is the concentration of the intermediate regulator *S*. The rate of expression of the intermediate regulator was assumed to be linearly dependent on 0A~P. For *spo0B, spo0E* and *rap* we assumed constant rates of expression (20).

In addition, the rate of expression of all genes in the model were also assumed to be proportional to the gene copy number and cell growth rate according to the following equation:

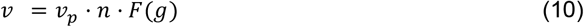

Where, *ν* represents the rate of gene expression, *ν*_*p*_ is given by *Eq. 9* with the specific expressions of each gene shown in Table S1, *n* is the gene copy number and *F*(*g*) is a function that relates cell size with the growth rate (*g*).

As in (20), the original proximal gene *spo0F* was assumed to be replicated at the start of DNA replication whereas the terminus proximal gene (*kinA*) was assumed to be replicated by the end of DNA replication, as illustrated in Figure 1B. The DNA replication period (*τ*_*rep*_) was assumed to be a function of the growth rate (*g*) with the following expression (20):

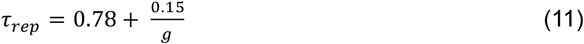

With the total cell cycle period (*τ* _*cyc*_) being given by (20):

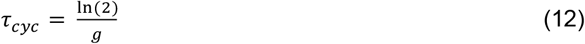

Meanwhile, *F*(*g*) follows the expression (20) :

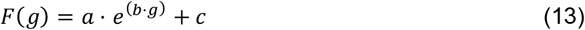

The values of *a, b*, and *c* were estimated from *in vivo* data in (20) and found to be *a* = 3.5, *b* = −ln (2), and *c* = 3.7.

### 4.2 Deterministic model of the biofilm matrix production network

To investigate the biofilm matrix production network assuming 0A~P pulsing dynamics we developed a model based on a previous mathematical model of the biofilm matrix production network (32). An illustration of the network is shown in Figure 1D. The model includes transcription, translation, and post-translation reactions of SinI, SinR, SlrR and TapA. The kinetic parameters and model reactions used can be found in Table S2.

The transcription and translation of *sinI, sinR, slrR*, and *tapA* are modelled in reactions R1-R4 of Table S2. Given that 0A~P induces the expression of *sinI*, the rate of production of SinI can be described according to the following equation:

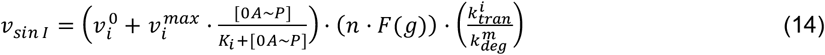

Where, transcription rate follows a Hill function with 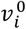 being the basal transcription rate and 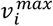 the maximum transcription rate when promoter is fully induced by 0A~P. The variable *K*_*i*_ is the half-maximal binding constant. The variable *n* corresponds to the average copy number and is estimated based on the following equation (15):

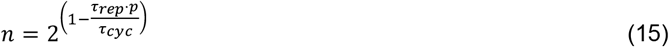

Where, *τ*_*rep*_ (DNA replication period) and *τ*_*cyc*_ (cell cycle length) are dependent on the growth rate as described by *Eqs. 11-12*, respectively. The variable *p* is the gene positioning relative to the *oriC*. As previously described, the function *F*(*g*) relates cell size with growth rate and was set to follow the same expression as in (15). The variables 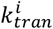 and 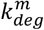 are the translation and RNA degradation rate, respectively.

We assume a constant rate of production for *sinR* (R2) since its expression is constitutive (45).

Meanwhile, given that the tetramer form of SinR (R6) acts as a repressor of *slrR* and *tapA*, similar to *Eq. 14*, the rate of production of SlrR (R3) and TapA (R4) can be modelled according to the following equations:

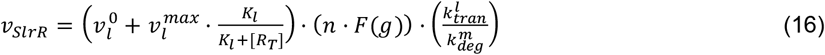

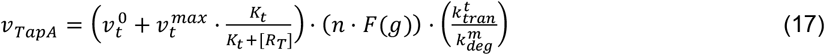

Where 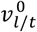 is the basal transcription rate, 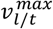 is the maximum transcription rate when the promoter is fully repressed by SinR tetramer form (*R*_*T*_). The variable *K*_*l*/*t*_ is the half-maximal binding constant. The subscripts *l*/*t* refer to *Eq*.*16 and 17*, respectively.

Moreover, SinR activity is regulated by SlrR and SinI due to complex formation. Namely, the SinI dimer (*I*_*d*_) interacts with the SinR dimer (*R*) and forms a SinI-SinR heterodimer (*IR*) (reaction R7) (46). On the other hand, SlrR dimer (*L*) associates with SinR dimer (*R*) and forms *LR* heterotetramer (R8). Notably, it is assumed that SinR (25) and SlrR are mostly dimeric (46).

The degradation rate of RNA 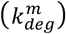 was set to 8.3 h^−1^ (47). The degradation rate of all proteins 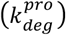 was set to 0.2 h^−1^ (48), except for SlrR which was set to 0.8 h^−1^, given that it is known to be an unstable protein (27). In addition, since growth rate (*g*) affects protein dilution the effective protein degradation rate 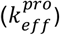 was assumed to be given by:

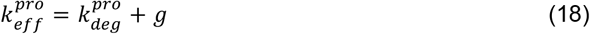

As in (32), the relative transcription, translation and *K*_*i*_ (*Eq. 14*) rates were set to ensure the bifurcation diagram resulted in the transitions from matrix OFF state to matrix ON state at a realistic growth rate and Spo0A~P level (Figure 4).

### 4.3 Stochastic model of the biofilm matrix production network

Given that biofilm matrix activation occurs stochastically (32), the deterministic biofilm matrix production model (Table S2) was adapted into a stochastic framework to explore whether pulsatile dynamics of 0A~P influence the probability of matrix activation. Unlike the deterministic model, the stochastic model explicitly incorporates transcription and translation reactions for each species. This approach accounts for downstream effects of random bursts in transcription and translation, which are critical when evaluating matrix activation. Additionally, we modeled the binding and unbinding dynamics of 0A~P to the promoter. This ensures that, under pulsatile 0A~P dynamics, the probability of promoter binding does not immediately follow the changes in 0A~P levels such that the time to reach the equilibrium probability of 0A~P binding to the promoter is accounted for. For the same reason, the binding and unbinding of the SinR tetramer to the *slrR* and *tapA* promoters is also explicitly modeled. Further, the transcription and translation rates for each mRNA were set to get a noise level in stochastic simulations to enable the stochastic activation and deactivation of biofilm matrix.

### 4.4 Model simulations

Deterministic model simulations (Figure 2, Figures 4-6) were done using the *ode15s* solver of MATLAB R2023b. The stochastic model was implemented in MATLAB R2023b, and the stochastic simulations (Figure 3) were done using the *SimBiology* tool of MATLAB R2023b.

In Figure 5, to investigate the role of DNA replication and post-replication period in deactivating biofilm matrix, *Eq. 11* was not used in the phosphorelay network to determine the DNA replication period and generate the corresponding 0A~P pulsing signal. Instead, model simulations were performed at an initial growth rate equal to 0.2 h^−1^. This corresponds to a DNA replication period equal to 1.53 h and post-replication period equal to 1.94 h (*τ*_*cyc*_ equal to 3.47 h). Next, DNA replication period was increased by increments of 0.1 h while post-replication period was kept constant (Figure 5A). At each increment we generated the corresponding 0A~P pulsing signal and ran a deterministic simulation of the biofilm matrix production network until the cell-cycle period leading to matrix deactivation was found (dark blue bar in Figure 5C). To generate Figure 5B the opposite was done. Post-replication period was increased by increments of 0.1 h while the DNA replication period was kept constant.

To generate Figure 6, we simulated the biofilm matrix production network for five different conditions, each differing in the inclusion or exclusion of growth rate effects on gene dosage, protein dilution, and/or 0A~P pulsing. Specifically, assuming an initial condition for which biofilm matrix is active (set to be at a growth rate equal to 0.2 h^−1^), we used the complete biofilm matrix production model with 0A~P pulsing as input to calculate the cell cycle length at which biofilm matrix production is deactivated. For this, we ran deterministic simulations for increasing cell cycle periods until reaching stead-state concentration of the model species (>100 h). Next, we determined what is the minimum period of the cell cycle leading to biofilm matrix deactivation.

To examine the impact of growth rate effects on protein dilution in relation to the deactivation cell cycle length, we modified the effective protein degradation rate to remove its dependence on growth rate. Consequently, *Eq. 19* was reformulated as:

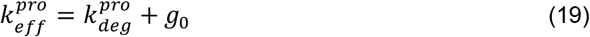

Here, *g*_0_ was set to 0.2 h^−1^.

Similarly, to examine the impact of gene dosage growth rate effects on the deactivation cell cycle length, we modified the gene average copy number equation (*Eq. 15*) to not be dependent on the growth rate. As such, *Eq. 15* was reformulated as:

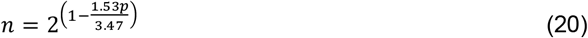

With 1.53 and 3.47 are the DNA replication time (*τ*_*rep*_) and the cell cycle length (*τ*_*cyc*_) for growth rate equal to 0.2 h^−1^, in accordance with *Eqs. 11*-*12*.

Finally, to assess the impact of 0A~P pulsing, we replaced the pulsatile 0A~P signal at a growth rate of 0.2 h^−1^ with a constant 0A~P signal equivalent to the average value of the pulsatile signal.

Combined effects (e.g. deleting growth rate effects on protein dilution and gene dosage) were investigated by applying the above modifications simultaneously in the network.

### 4.6 Model fitting results

To estimate the biofilm matrix activation and deactivation rate (*k*_*ON*_ and *k*_*OFF*_, respectively) we simultaneously fit *Eq. 3* to ‘*Initially OFF*’ cells and *Eq. 4* to ‘*Initially ON*’ cells. We constrained the fitting such that *k*_*ON*_ rate is set to be same between constant and a pulsatile 0A~P input. The variable *k*_*OFF*_ was set to be a free parameter. To find the best fitting rates constants, we defined an objective function for optimization. This function minimizes the sum of the squared differences between the stochastic data (light blue and orange lines in Figure 3D) and predictions. Optimization was done using *fmincon* function of MATLAB R2023b.

To estimate the biofilm deactivation threshold when 0A~P oscillation period and cell cycle period are the same (Figure 4D) we applied a power-law fit (*Eq. 21)* using the non-linear least square method.

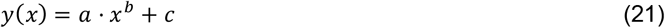

The fitting results predict *a* = 0.06, *b* = 2.59, *c* = 0.14. The fitting R^2^ value is 0.99.

## 5. Data availability statement

A data software package with the deterministic and stochastic models used, and all figure generation scripts is deposited in GitHub (https://github.com/PalmaCristina/biofilm-matrix-deactivation).

## 6. Supplemental data

Supplemental data available online.

## 7. Author contributions

*Author contributions*: O.A.I and J.J.T conceived and supervised the study. C.S.D.P. executed the model implementation, simulation and results interpretation, to which D.J.H contributed to. C.S.D.P. drafted all documents which were revised by all co-authors.

## 8. Acknowledgements

This work was supported by National Science Foundation [MCB-2204402 to O.A.I (co-PI) and J.J.T. (PI)], by the Jenny and Antti Wihuri Foundation [to C.S.D.P.] and by the National Science Foundation Graduate Research Fellowship [1842494 to D.J.H].

## SUPPLEMENTAL MATERIAL FOR

## Supplementary Tables

**Table S1:**
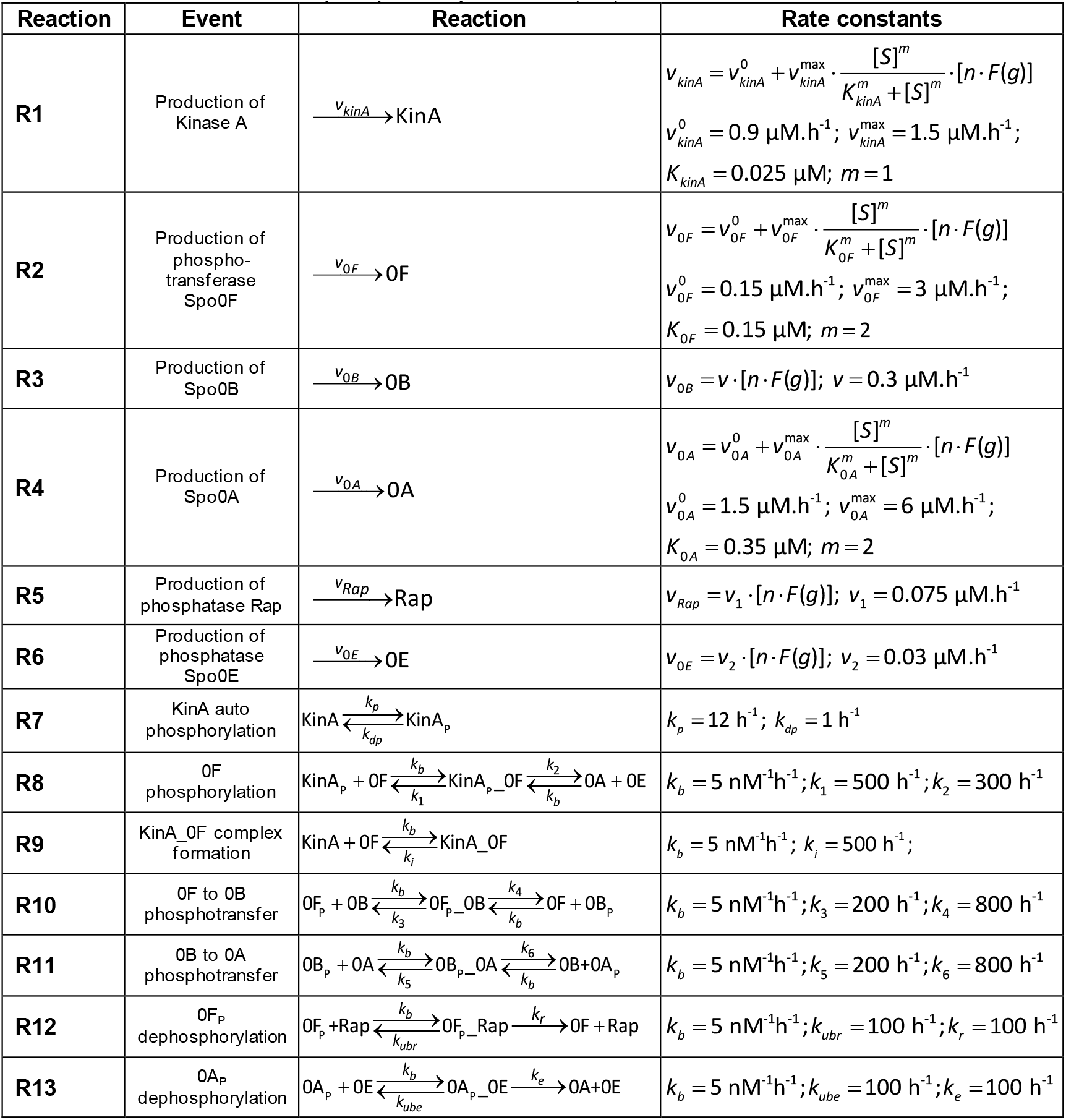
Kinetic parameters and model reactions of the phosphorelay model (Methods section 4.1). Model based on the phosphorelay model in (1). Subscript *P* marks phosphorylated forms of proteins. The underscore symbol denotes protein complexes. Reaction R12-R13 were assumed to have catalytic rate constants 100 hr^−1^ (2). Protein degradation reactions are not included in the table, for all proteins the degradation rate was fixed at 0.3 hr^−1^ (1). Effective protein degradation rate is dependent on the growth rate as described by *Eq. 17*. In the table, *g* stands for ‘growth rate’, *n* for the copy number of the gene, and *F(g)* is the function describing how the cell volume changes with the growth rate (Methods Section 4.1). All other rate constants were estimated from the *in vitro* measurements of phosphorelay kinetics (1,3).

| Reaction | Event | Reaction | Rate constants |
| --- | --- | --- | --- |
| R1 | Production of Kinase A | $\xrightarrow{v_{kinA}} \text{KinA}$ | $v_{kinA} = v_{kinA}^0 + v_{kinA}^{\max} \cdot \frac{[S]^m}{K_{kinA}^m + [S]^m} \cdot [n \cdot F(g)]$<br>$v_{kinA}^0 = 0.9 \mu\text{M} \cdot \text{h}^{-1}$; $v_{kinA}^{\max} = 1.5 \mu\text{M} \cdot \text{h}^{-1}$;<br>$K_{kinA} = 0.025 \mu\text{M}$; $m = 1$ |
| R2 | Production of phospho-transferase Spo0F | $\xrightarrow{v_{0F}} \text{0F}$ | $v_{0F} = v_{0F}^0 + v_{0F}^{\max} \cdot \frac{[S]^m}{K_{0F}^m + [S]^m} \cdot [n \cdot F(g)]$<br>$v_{0F}^0 = 0.15 \mu\text{M} \cdot \text{h}^{-1}$; $v_{0F}^{\max} = 3 \mu\text{M} \cdot \text{h}^{-1}$;<br>$K_{0F} = 0.15 \mu\text{M}$; $m = 2$ |
| R3 | Production of Spo0B | $\xrightarrow{v_{0B}} \text{0B}$ | $v_{0B} = v \cdot [n \cdot F(g)]$; $v = 0.3 \mu\text{M} \cdot \text{h}^{-1}$ |
| R4 | Production of Spo0A | $\xrightarrow{v_{0A}} \text{0A}$ | $v_{0A} = v_{0A}^0 + v_{0A}^{\max} \cdot \frac{[S]^m}{K_{0A}^m + [S]^m} \cdot [n \cdot F(g)]$<br>$v_{0A}^0 = 1.5 \mu\text{M} \cdot \text{h}^{-1}$; $v_{0A}^{\max} = 6 \mu\text{M} \cdot \text{h}^{-1}$;<br>$K_{0A} = 0.35 \mu\text{M}$; $m = 2$ |
| R5 | Production of phosphatase Rap | $\xrightarrow{v_{Rap}} \text{Rap}$ | $v_{Rap} = v_1 \cdot [n \cdot F(g)]$; $v_1 = 0.075 \mu\text{M} \cdot \text{h}^{-1}$ |
| R6 | Production of phosphatase Spo0E | $\xrightarrow{v_{0E}} \text{0E}$ | $v_{0E} = v_2 \cdot [n \cdot F(g)]$; $v_2 = 0.03 \mu\text{M} \cdot \text{h}^{-1}$ |
| R7 | KinA auto phosphorylation | $\text{KinA} \xrightleftharpoons[k_{dp}]{k_p} \text{KinA}_p$ | $k_p = 12 \text{ h}^{-1}$; $k_{dp} = 1 \text{ h}^{-1}$ |
| R8 | 0F phosphorylation | $\text{KinA}_p + \text{0F} \xrightleftharpoons[k_1]{k_b} \text{KinA}_p\text{-0F} \xrightleftharpoons[k_b]{k_2} \text{0A} + \text{0E}$ | $k_b = 5 \text{ nM}^{-1} \text{h}^{-1}$; $k_1 = 500 \text{ h}^{-1}$; $k_2 = 300 \text{ h}^{-1}$ |
| R9 | KinA_0F complex formation | $\text{KinA} + \text{0F} \xrightleftharpoons[k_i]{k_b} \text{KinA\_0F}$ | $k_b = 5 \text{ nM}^{-1} \text{h}^{-1}$; $k_i = 500 \text{ h}^{-1}$; |
| R10 | 0F to 0B phosphotransfer | $\text{0F}_p + \text{0B} \xrightleftharpoons[k_3]{k_b} \text{0F}_p\text{-0B} \xrightleftharpoons[k_b]{k_4} \text{0F} + \text{0B}_p$ | $k_b = 5 \text{ nM}^{-1} \text{h}^{-1}$; $k_3 = 200 \text{ h}^{-1}$; $k_4 = 800 \text{ h}^{-1}$ |
| R11 | 0B to 0A phosphotransfer | $\text{0B}_p + \text{0A} \xrightleftharpoons[k_5]{k_b} \text{0B}_p\text{-0A} \xrightleftharpoons[k_b]{k_6} \text{0B} + \text{0A}_p$ | $k_b = 5 \text{ nM}^{-1} \text{h}^{-1}$; $k_5 = 200 \text{ h}^{-1}$; $k_6 = 800 \text{ h}^{-1}$ |
| R12 | 0F<sub>p</sub> dephosphorylation | $\text{0F}_p + \text{Rap} \xrightleftharpoons[k_{ubr}]{k_b} \text{0F}_p\text{-Rap} \xrightarrow{k_r} \text{0F} + \text{Rap}$ | $k_b = 5 \text{ nM}^{-1} \text{h}^{-1}$; $k_{ubr} = 100 \text{ h}^{-1}$; $k_r = 100 \text{ h}^{-1}$ |
| R13 | 0A<sub>p</sub> dephosphorylation | $\text{0A}_p + \text{0E} \xrightleftharpoons[k_{ube}]{k_b} \text{0A}_p\text{-0E} \xrightarrow{k_e} \text{0A} + \text{0E}$ | $k_b = 5 \text{ nM}^{-1} \text{h}^{-1}$; $k_{ube} = 100 \text{ h}^{-1}$; $k_e = 100 \text{ h}^{-1}$ |

**Table S2:** Kinetic parameters and model reactions of the biofilm matrix production deterministic model. The rate constants were converted from (4) stochastic model, assuming a cell volume of 4fL (5). In the table, *g* stands for ‘growth rate’, *n* for the copy number of the gene, and *F(g)* is the function describing how the cell volume changes with the growth rate (Methods Section 4.2). The degradation rate of RNA 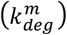 was set to 8.3 h^−1^ (6). The degradation rate of all proteins 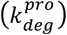 was set to 0.2 h^−1^ (7), except for SlrR which was set to 0.8 h^−1^, given that it is known to be an unstable protein (8). The relative transcription, translation and *K*_*i*_ rates were set to ensure the bifurcation diagram resulted in the transitions from matrix OFF state to matrix ON state at a realistic growth rate and 0A~P level.

| Reaction | Event | Reaction | Rate constants |
| --- | --- | --- | --- |
| **R1** | Production of SinI | $\xrightarrow{v_{sinI}} I$ | $v_{sinI} = v_i^0 + v_i^{\max} \cdot \frac{[0A \sim P]}{K_i + [0A \sim P]} \cdot [n \cdot F(g)] \cdot \frac{k_{tran}^i}{k_{deg}^m}$<br>$v_i^0 = 0; v_i^{\max} = 0.03 \mu\text{M} \cdot \text{h}^{-1}; K_i = 0.01 \mu\text{M};$<br>$k_{tran}^i = 400 \text{ h}^{-1}; k_{deg}^m = 8.3 \text{ h}^{-1};$ |
| **R2** | Production of SinR | $\xrightarrow{v_{sinR}} R$ | $v_{sinR} = v_r \cdot [n \cdot F(g)] \cdot \frac{k_{tran}^r}{k_{deg}^m}$<br>$v_r = 0.05 \mu\text{M} \cdot \text{h}^{-1}; k_{tran}^r = 200 \text{ h}^{-1}; k_{deg}^m = 8.3 \text{ h}^{-1}$ |
| **R3** | Production of SlrR | $\xrightarrow{v_{slrR}} L$ | $v_{slrR} = v_l^0 + v_l^{\max} \cdot \frac{K_l}{K_l + [R_T]} \cdot [n \cdot F(g)] \cdot \frac{k_{tran}^l}{k_{deg}^m}$<br>$v_l^0 = 0; v_l^{\max} = 0.046 \mu\text{M} \cdot \text{h}^{-1}; K_l = 0.9 \text{ nM};$<br>$k_{tran}^l = 200 \text{ h}^{-1}; k_{deg}^m = 8.3 \text{ h}^{-1};$ |
| **R4** | Production of TapA | $\xrightarrow{v_{tapA}} T$ | $v_{tapA} = v_t^0 + v_t^{\max} \cdot \frac{K_t}{K_t + [R_T]} \cdot [n \cdot F(g)] \cdot \frac{k_{tran}^t}{k_{deg}^m}$<br>$v_t^0 = 0; v_t^{\max} = 0.01 \mu\text{M} \cdot \text{h}^{-1}; K_t = 2.1 \text{ nM};$<br>$k_{tran}^t = 200 \text{ h}^{-1}; k_{deg}^m = 8.3 \text{ h}^{-1};$ |
| **R5** | Formation of SinI dimer | $I + I \xrightleftharpoons[k_{udi}]{k_{di}} I_d$ | $k_{di} = 722 \mu\text{M}^{-1} \cdot \text{h}^{-1}; k_{udi} = 169 \text{ h}^{-1};$ |
| **R6** | Formation of SinR tetramer | $R + R \xrightleftharpoons[k_{udr}]{k_{dr}} R_T$ | $k_{dr} = 722 \mu\text{M}^{-1} \cdot \text{h}^{-1}; k_{udr} = 247 \text{ h}^{-1};$ |
| **R7** | Formation of SinI-SinR heterodimer | $I_d + R \xrightarrow{k_{b1}} 2IR$ | $k_{b1} = 770 \mu\text{M}^{-1} \cdot \text{h}^{-1}$ |
| **R8** | Formation of SlrR2-SinR2 heterotetramer | $L + R \xrightleftharpoons[k_{dlr}]{k_{b2}} LR$ | $k_{b2} = 770 \mu\text{M}^{-1} \cdot \text{h}^{-1}; k_{dlr} = 0.99 \text{ h}^{-1}$ |

## Supplementary Figures

**Figure S1:**
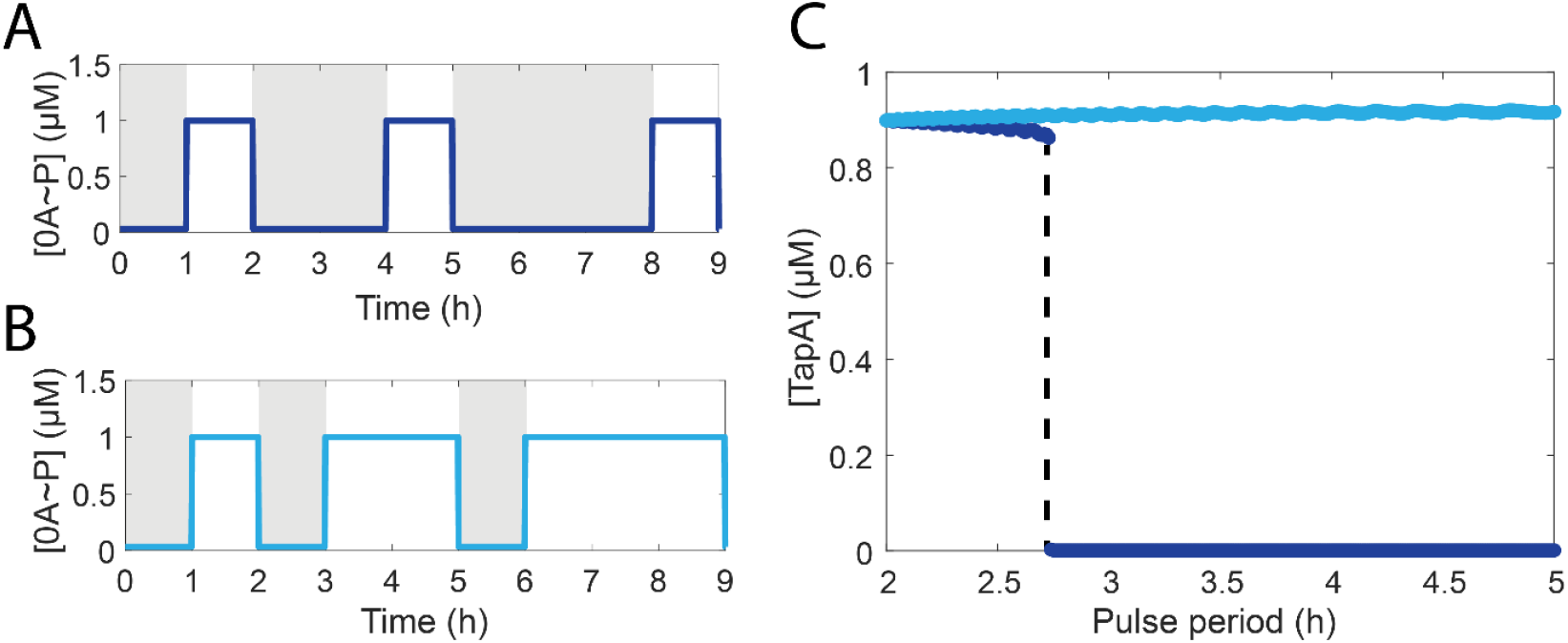
Effects of 0A~P OFF and ON time in matrix deactivation. **(A)** 0A~P square pulse dynamics for increasing OFF time length (grey area) and constant ON time (white area). **(B)** 0A~P square pulse dynamics for constant OFF time length (grey area) and increasing ON time (white area). The initial pulse given was the same with an ON and OFF time-length equal to 1 h. Square pulse height was kept constant and equal to 1. **(C)** Dark blue line is the result for the pulse dynamics shown in A, i.e., ON time is kept constant while OFF time increases. Light blue line is the result for the pulse dynamics shown in (B), i.e., OFF time is kept constant while ON time increases.

**Figure S2:**
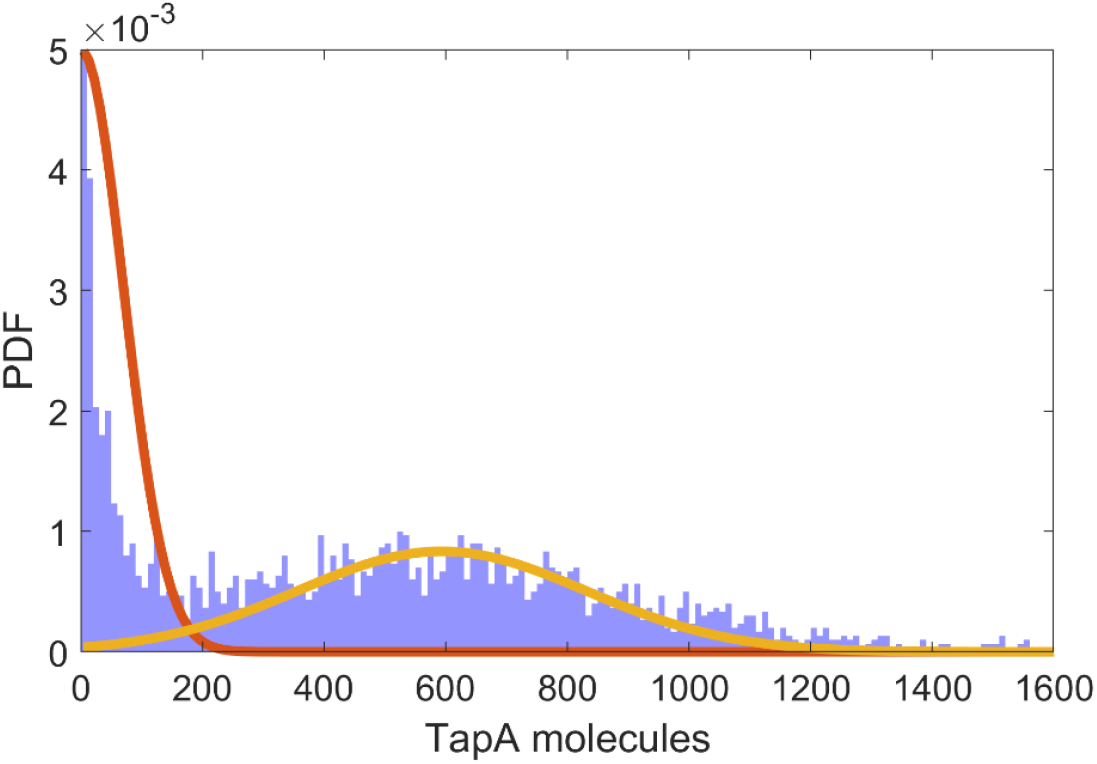
Histogram of TapA molecules obtained from 3000 stochastic simulations (blue bars). Simulation time was set to 50 h. Solid lines represent Gaussian fits derived from a Gaussian Mixture Model with 2 components fitted to the data. The value of 200 molecules was set to be the threshold to distinguish between a biofilm matrix production active and inactive.

